# Uterine histotroph and conceptus development. III. Adrenomedullin stimulates proliferation, migration and adhesion of porcine trophectoderm cells via AKT-TSC2-MTOR cell signaling pathway

**DOI:** 10.1101/2022.10.10.511652

**Authors:** Bangmin Liu, Sudikshya Paudel, William L Flowers, Jorge A Piedrahita, Xiaoqiu Wang

## Abstract

Adrenomedullin (ADM) as a highly conserved peptide hormone has been reported to increase significantly in the uterine lumen during the peri-implantation period of pregnancy in pigs, but its functional roles in growth and development of porcine conceptus (embryonic/fetus and its extra-embryonic membranes) as well as underlying mechanisms remain largely unknown. Therefore, we conducted *in vitro* experiments using our established porcine trophectoderm cell line (pTr1) isolated from Day-12 porcine conceptuses to test the hypothesis that porcine ADM stimulates cell proliferation, migration and adhesion via AKT-TSC2-MTOR cell signaling pathway in pTr1 cells. Porcine ADM at 10^-7^ M stimulated (*P*<0.05) pTr1 cell proliferation, migration and adhesion by 1.4-, 1.5- and 1.2-folds, respectively. These ADM-induced effects were abrogated (*P*<0.05) by siRNA-mediated knockdown of ADM (siADM) and its shared receptor component calcitonin-receptor-like receptor (CALCRL; siCALCRL), as well as by rapamycin, the inhibitor of mechanistic target of rapamycin (MTOR). Using siRNA mediated knockdown of CALCRL coupled with Western blot analyses, ADM signaling transduction was determined in which ADM binds to CALCRL to increase phosphorylation of MTOR, its downstream effectors (4EBP1, P70S6K, and S6), and upstream regulators (AKT and TSC2). Collectively, these results suggest that porcine ADM in histotroph act on its receptor component CALCRL to activate AKT-TSC2-MTOR, particularly MTORC1 signaling cascade, leading to elongation,migration and attachment of conceptuses.

## INTRODUCTION

In all mammalian species, the greatest constraint to reproductive performance is embryonic mortality, which counts 20%-30% of embryos in most cases; however, the true rate of early pregnancy loss is close to 50% due to the high number of pregnancies that are not recognized in the first 2 to 4 weeks after conception (Bazer and First 1983; Patel and Lessey 2011; Racowsky 2002). Intrauterine growth restriction (IUGR), defined as impaired growth and development of fetuses and their organs during pregnancy, is also commonly observed in mammals, and in turn leads to “runt” offspring with poor survival in the neonatal period of life (McMillen and Robinson 2005; Wu et al. 2006; Bauer et al. 1998; Wang et al. 2008; Wang et al. 2009; Wang et al. 2010; Liu et al. 2013; Wang et al. 2013; Wu et al. 2013; Wang et al. 2014c; Wang et al. 2018). Of particular interests, pigs exhibit a high incidence of early embryonic death (30-40%), and the most severe naturally occurring IUGR (15-25%) (Wu et al. 2008; Wu et al. 2006; Redmer et al. 2004; Wang et al. 2010; Wang et al. 2014c; Vallet et al. 2011; Vallet et al. 2014; Mesa et al. 2006; Tuchscherer et al. 2000; Leenhouwers et al. 2001; Wang et al. 2017; Tarraf and Knight 1995a, b; Chen and Dziuk 1993; Vallet et al. 1994; Vallet and Christenson 1993; Vallet et al. 2013). This is likely due to the asynchrony between uterine and conceptus (embryo/fetus and its extra-embryonic membranes) development that limits surface area of placentation at the maternal-conceptus interface, compromises conceptus development and results in fetal crowding, i.e., limited uterine capacity during gestation (Geisert and Schmitt 2002; Pope 1994; Town et al. 2005; Dziuk 1968; Bazer et al. 1969b; Bazer et al. 1969a). However, the underlying mechanisms by which conceptuses grow and develop in concert with intrauterine environment are poorly understood.

During the peri-implantation period of pregnancy, porcine conceptuses undergo rapid elongation from spherical to tubular and filamentous forms that requires proliferation, migration, and cytoskeleton reorganization of trophectoderm (Tr) cells (Bazer 1975; Spencer and Bazer 2004). This orchestrated process is highly dependent on the composition of histotroph (Bazer et al. 2014; Kong et al. 2014; Wang et al. 2014a; Wang et al. 2014b; Wang et al. 2014d; Bazer et al. 2015; Wang et al. 2015a; Wang et al. 2015b; Lenis et al. 2016; Wang et al. 2016a; Wang et al. 2016b; Lenis et al. 2018; Gao et al. 2009e, d, c, b, a; Gao et al. 2009f; Kim et al. 2011a; Kim et al. 2011b; Bazer et al. 2012; Kim et al. 2011c), a complex mixture of molecules secreted or transported into the uterine lumen by uterine luminal (LE), superficial glandular (sGE), and deep glandular (dGE) epithelia as well as conceptus Tr (Roberts and Bazer 1988; Spencer et al. 2004; Bazer 1975). Recently, we reported that adrenomedullin (ADM; a highly conserved peptide in mammals) increases significantly in the uterine lumen, as does expression of ADM mRNA and protein, along with its receptor components calcitonin-receptor-like receptor (CALCRL), receptor activity-modifying protein 2 (RAMP2) and 3 (RAMP3) in uterine endometria and conceptuses during the peri-implantation period of pregnancy in pigs (Paudel et al. 2021). Using a well-established and well-studied porcine Tr cell line (pTr1 cells) that exhibits numerous properties of pTr cells *in vivo* (Kong et al. 2012; Kong et al. 2014; Ka et al. 2001; Jaeger et al. 2005), we also demonstrated that ADM stimulates proliferation and increases phosphorylation of mechanistic target of rapamycin (MTOR) in pTr1 cells (Paudel et al. 2021). However, in addition to proliferation, little is known about whether ADM affect migration and adhesion of pTr1 cells or the underlying mechanisms, if any, responsible for those effects. To achieve this goal, pTr1 cells were subjected to siRNA-mediated knockdown and treated with ADM to determine effects on proliferation, migration and adhesion, as well as expressions of total and phosphorylated proteins of MTOR, eukaryotic initiation factor (eIF) 4E-binding protein-1 (4EBP1), ribosomal protein S6 kinase 1 (P70S6K), ribosomal protein S6 (S6), tuberous sclerosis 2 protein (TSC2), and protein kinase B (AKT). Our results indicate that ADM acts on its receptor component CALCRL to increase cell proliferation, migration and adhesion; and that these stimulatory effects are mediated through at least activation of the AKT-TSC2-MTORC1 cell signaling cascade.

## MATERIAL AND METHODS

### Cell culture

An established spontaneously immortalized porcine trophectoderm primary cell line (pTr1; a kind gift from Dr. Robert C. Burghardt, Texas A&M University, College Station, TX) from Day-12 porcine conceptuses was developed, propagated, and used in the present *in vitro* studies as described previously (Ka et al. 2001; Jaeger et al. 2005; Kong et al. 2014; Paudel et al. 2021). The pTr1 cells exhibit numerous properties of pTr cells in *vivo* (Ka et al. 2001). The cells were cultured in complete medium (CM; Dulbecco modified Eagle medium/Nutrient Mixture F-12, DMEM/F-12; Gibco BRL, Grand Island, NY), with 10% fetal bovine serum (FBS; Gibco BRL), 50 U/ml penicillin, 50 μg/ml streptomycin, 0.1 mM each for nutritionally nonessential amino acids (NEAA), 1 mM sodium pyruvate, 2 mM glutamine, and 4 μg/ml insulin. The medium was replaced every 2 days. When the density of cells in the dishes reached about 80%confluence, subcultures of cells were prepared at a ratio of 1:3, and frozen stocks of cells were preserved at each passage. For the experiments, cultures of pTr1 cells (between passages 8 and 20) were grown in CM to 20%-30% confluences, serum- and insulin-starved for 24 h in customized medium, and then treated with porcine ADM (ADM; 4095741; Bachem Americas, Inc.,Torrance, CA, USA) at 10^-7^ M in basal medium (BM; DMEM/F-12 with 5% FBS and 1 ng/ml insulin). The experiments were replicated three times independently.

### Small-interfering RNA knockdown

The pTr1 cells were transfected with 60 nM nontargeting control siRNA (siNTC; D-001810-01-05), siRNA targeting porcine *ADM* (siADM; CTM-626827) or siRNA targeting porcine *CALCRL* (siCALCRL; CTM-626822) (ON-TARGETplus SMARTpool; Dharmacon, Lafayette, CO). Transfection was performed using Lipofectamine 2000 diluted (0.4% v/v; Invitrogen) in Opti-MEM reduced serum medium containing 2% charcoal stripped FBS (SFBS; Gibco BRL) according to the manufacturer’s instructions. After 48 h transfection, the medium was removed, and the pTr1 cells were subcultured in the BM with or without 10^-7^ M ADM as described previously (Paudel et al. 2021).

### Cell proliferation assay

The pTr1 cells were cultured (1×10^4^ cells/0.4 ml/well) in 24-well plates (Costar no. 3524; Corning, Corning, NY, USA) in complete medium (CM) until the monolayer reached 30% confluence and then subject to siRNA-mediated knockdown. After 48 h transfection, cells (n=6 wells per treatment) were subcultured in 0.4 ml BM with or without 10^-7^ M ADM for additional 48 h. Cell numbers were determined after 48 of incubation as described previously (Raspotnig et al. 1999; Kong et al. 2014; Wang et al. 2015a; Wang et al. 2015b; Paudel et al. 2021). Briefly, medium was removed from cells by vacuum aspiration, cells were fixed in 50% ethanol for 30 min, and fixative was removed by vacuum aspiration. Fixed cells were stained with Janus Green B in PBS (pH 7.2) for 3 min at room temperature. The stain was immediately removed using a vacuum aspirator, and the whole plate was sequentially dipped into water and destained by gentle shaking. The remaining water was removed by shaking, after which stained cells were immediately lysed in 0.5 N HCl and absorbance readings were taken at 595 nm using a microplate reader. As described previously (Raspotnig et al. 1999), cell numbers were calculated from absorbance readings using the following formula: cell number = (absorbance - 0.00462)/0.00006926.

### Cell migration assay

Cell migration was determined by a 12-h wound-healing assay using 6-well plates as described previously (Liu et al. 2019). Briefly, the pTr1 cells (2×10^5^ cells/2 ml/well) were seeded in 6-well plates (83.3920; Sarstedt, Nümbrecht, Germany) in CM until the monolayer reached 80-90% confluence, and subject to siRNA-mediated knockdown (n=6 wells per treatment). After 48 h transfection, a cell-free area was created by a straight scratch to the center of each well using a 200-μl pipette tip. The off-scratched cells were removed using a vacuum aspirator. After rinsed with PBS (pH 7.2) twice, the remaining cells were subcultured in 2 ml BM with or without 10^-7^ M ADM for additional 12 h. For migration assay with MTOR inhibitor, pTr1 cells were pre-treated with 50 nM rapamycin (R8781-200UL; Sigma-Aldrich) for 6 h before applying straight scratch, and then treated with or without 10^-7^ M ADM. The images of each well at 0 and 12 h post scratching was captured by EVOSTM M5000 Imaging system (Thermo Fisher Scientific) and the area of cell migration measured using Image J (NIH).

### Cell adhesion assay

Cell adhesion assays were performed as described previously (Wang et al. 2016a). Briefly, the pTr1 cells (5×10^5^ cells/5 ml/flask) were seeded in T25 flasks (83.3910.002; Sarstedt) in CM. When the monolayer reached 80-90% confluence, pTr1 cells were subject to siRNA-mediated knockdown. After 48 h transfection, cells were re-seeded (1×10^4^ cells/0.2 ml/well; n=6 wells per treatment) in 96-well plates, and allowed to attach for 2 h (37 °C, 5% CO2) with or without the presence of 10^-7^ M ADM. After 2 h for attachment, nonadherent cells were removed with medium by vacuum aspiration and rinsed twice in PBS. The wells were fixed in 10% v/v formalin in PBS. Cell numbers were determined as described in proliferation assay. For adhesion assay with MTORC1 inhibitor, pTr1 cells were pre-treated with 50 nM rapamycin in T25 flasks for 6 h before re-seeding in 96-well plates.

### Protein extraction and western blot analysis

The pTr1 cell (2×10^5^ cells/2 ml/well) were seeded in 6-well plates in CM until the monolayer reached 80-90% confluence, and subject to siRNA-mediated knockdown. After 48 h transfection, cells were subcultured in 2 ml BM with or without 10^-7^ M ADM for additional 30- and 60-min. Cells in each well were then lysed for 30 min on ice in 0.1 ml of a lysis buffer consisting of 1% Triton X-100, 0.5% Nonidet P-40, 150 mM NaCl, 10 mM Tris-HCl (pH 7.6), 1 mM EDTA, 1 mM ethylene glycol tetra-acetic acid, 0.2 mM Na_3_VO_4_, 0.2 mM phenylmethylsulfonylfluoride, 50 mM NaF, 30 mM Na_4_P_2_O_7_, and 1% protease inhibitor cocktail (Kong et al. 2014). The cell lysates were passed through a sterile 26-gauge needle and centrifuged (21000 × g for 15 min at 4 °C). Protein concentration in the supernatant was determined using the Bradford protein assay with bovine serum albumin at the standard (Wang et al. 2015a). Western blot analysis was performed as described previously (Cummings et al. 2022). Extracted protein (28 μg/sample) were denatured at 95°C for 10 min, separated using sodium dodecyl sulphate polyacrylamide gel electrophoresis (SDS-PAGE; 12% gel at 70 V for 30 min to pass through stacker gel, and 110 V for additional 2.5-3 h) and transferred to a nitrocellulose membrane overnight (~16 h) at 15 V using the Bio-Rad Transblot (Bio-Rad). Membranes were blocked in 5% fat-free milk in Tris buffered saline with Tween-20 (TBST; 20 mM Tris, 150 mM NaCl, and 0.1 % Tween-20, pH 7.5) for 3 h and then incubated with a primary antibody detecting total (t-) and phosphorylated (p-) MTOR, 4EBP1, P70S6K, S6, TSC2 and AKT at 4°C overnight (~16 h) with gentle rocking. After washing three times (10 min per time) with TBST, the membranes were incubated for 2 h with a secondary antibody. The membranes were then washed with TBST, followed by development using enhanced chemiluminescence detection (1705060, Bio-Rad) according to the manufacturer’s instructions. Rabbit anti-glyceraldehyde-3-phosphate dehydrogenase (GAPDH) polyclonal immunoglobulin G was used as loading control antibody for each blot. Western blots were analyzed by measuring the intensity of light emitted from correctly sized bands under ultraviolet light using a ChemiDocTM MP Imaging System (Bio-Rad) and Image Lab software (version 6.1; Bio-Rad). Multiple exposures of each Western blot were performed to ensure the linearity of chemiluminescence signals. Information for all antibodies is listed in **Table 1**.

**Table 1.** Antibodies and dilutions used for Western blot analysis.

| Antibody | Catalog No. | Dilution | Source |
| --- | --- | --- | --- |
| Primary antibody<sup>a</sup> |  |  |  |
| t-MTOR | 2972 | 1:1000 | Rabbit |
| p-MTOR (Ser2448) | 2971 | 1:1000 | Rabbit |
| t-4EBP1 | 9452 | 1:1000 | Rabbit |
| p-4EBP1 (Thr70) | 9455 | 1:1000 | Rabbit |
| t-P70S6K | 9202 | 1:1000 | Rabbit |
| p-P70S6K (Th389) | 9206 | 1:1000 | Mouse |
| t-S6 | 2317 | 1:1000 | Mouse |
| p-S6 (Ser240/244) | 2215 | 1:1000 | Rabbit |
| t-TSC2 | 4308 | 1:1000 | Rabbit |
| p-TSC2 (Ser939) | 3615 | 1:1000 | Rabbit |
| t-AKT | 9272 | 1:1000 | Rabbit |
| p-AKT (Thr308) | 9275 | 1:1000 | Rabbit |
| GAPDH | 2118 | 1:2000 | Rabbit |
| Secondary antibody<sup>b</sup> |  |  |  |
| Anti-Mouse IgG Antibody (H+L), Peroxidase<sup>b</sup> | PI-2000 | 1:2000 | Horse |
| Anti-Rabbit IgG Antibody (H+L), Peroxidase<sup>b</sup> | PI-1000 | 1:2000 | Goat |

<sup>a</sup> Antibodies purchased from Cell Signaling Technology
<sup>b</sup> Antibodies from Vector Laboratories

### Statistical analysis

Normality of data and homogeneity of variance were tested using the Shapiro-Wilk test and Brown-Forsythe test in Statistical Analysis System, respectively (version 8.1; SAS Institute). Data were analyzed by least squares one-way analysis of variance (ANOVA) and post hoc analysis (the Fisher least significant difference) with each well/band as an experimental unit. All analyses were performed using SAS. Data are expressed as means with SEM. *P*<0.05 was considered statistically significant.

## RESULTS

### ADM stimulates proliferation of pTr1 cells via its receptor component CALCRL

We first confirmed the effects of ADM on pTr1 cell proliferation using siRNA-mediated knockdown of endogenous ADM (siADM) and its receptor component CALCRL (siCALCRL; **Figure 1**). Compared with siNTC control, exogenous ADM at 10^-7^ M increased (*P*<0.05) pTr1 cell proliferation by 1.4-fold in siNTC+ADM group at 48 h. On the other hand, inhibition of endogenous ADM by siADM decreased (*P*<0.05) pTr1 cell proliferation by 33% and 29% in the absence (siADM *versus* siNTC) and presence (siADM+ADM *versus* siNTC+ADM) of exogenous ADM, respectively. When endogenous ADM was inhibited by siADM, exogenous ADM at 10^-7^ M simulated (*P*<0.05) pTr1 cell proliferation by 1.5-fold at 48 h (siADM+ADM *versus* siADM). Moreover, inhibition of CALCRL, the shared component of ADM receptors, abrogated the ADM-derived proliferation of pTr1 cells by 41% in siCALCRL+ADM group as compared to siNTC+ADM control.

**Figure 1.**
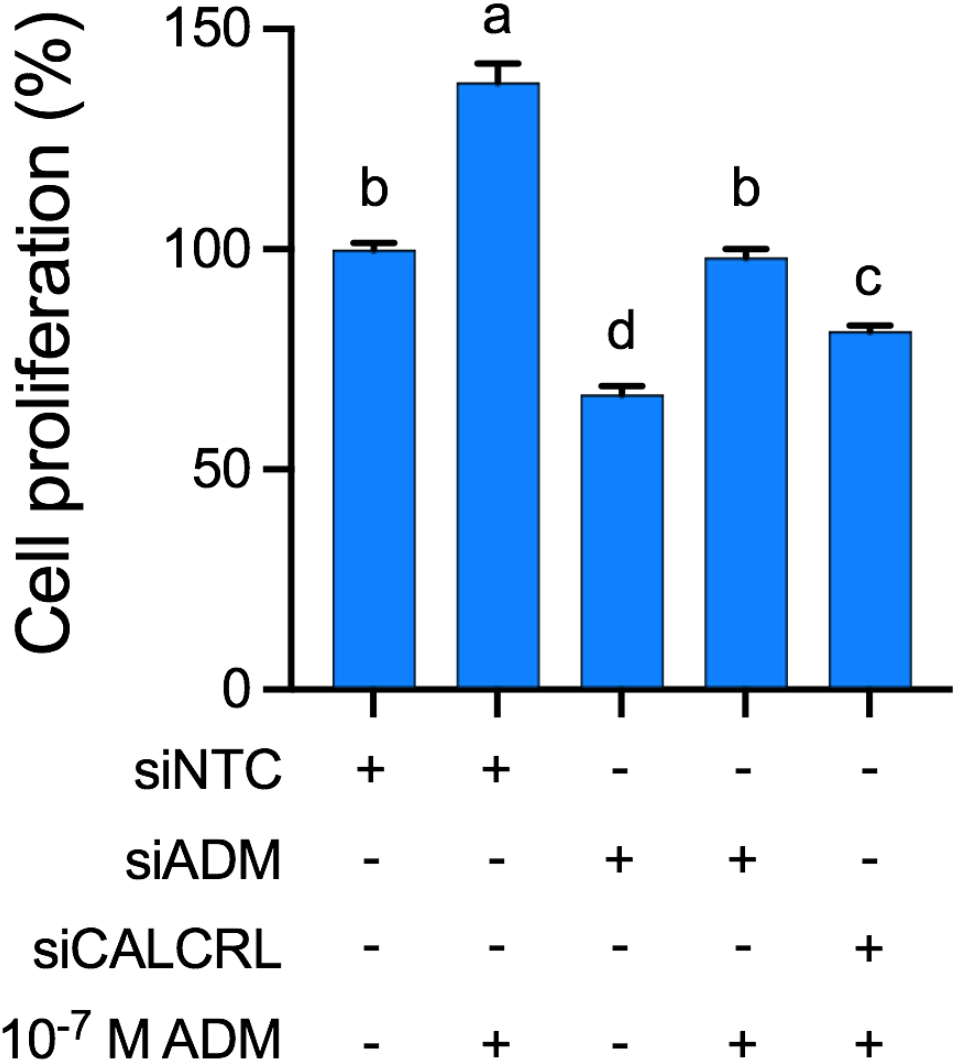
ADM stimulates proliferation of pTr1 cells via its receptor component CALCRL. The pTr1 cells (n=6 wells) were transfected with siNTC, siADM or siCALCRL for 48 h, and then subjected to ADM treatment at 10 M. Cell numbers were determined after additional 48 h of incubation. Data are expressed as a percentage relative to siNTC treated cells. siNTC, nontargeting control siRNA; siADM, siRNA targeting porcine *ADM*; siCALCRL, siRNA targeting porcine *CALCRL*. Different superscript letters denote significant differences (*P*<0.05) among groups. Data are presented as means and SEM.

### ADM stimulates migration of pTr1 cells via its receptor component CALCRL

Next, we determined the effects of ADM on migration of pTr1 cells using a 12-h wound-healing assay coupled with siRNA-mediated knockdown (**Figure 2**). After 48 h transfection and additional 12 h ADM treatment, exogenous ADM at 10^-7^ M increased (*P*<0.05) pTr1 cell migration by 1.5-fold in siNTC+ADM group as compared to siNTC control. Inhibition of endogenous ADM by siADM decreased (*P*<0.05) pTr1 cell migration by 46% and 23% in the absence (siADM *versus* siNTC) and presence (siADM+ADM *versus* siNTC+ADM) of exogenous ADM, respectively. When endogenous ADM was inhibited by siADM, exogenous ADM at 10^-7^ M simulated (*P*<0.05) pTr1 cell migration by 2.1-fold at 12 h (siADM+ADM *versus* siADM). In addition, inhibition of CALCRL abrogated (*P*<0.05) the ADM-derived migration of pTr1 cells by 42% in siCALCRL+ADM group as compared to siNTC+ADM control.

**Figure 2.**
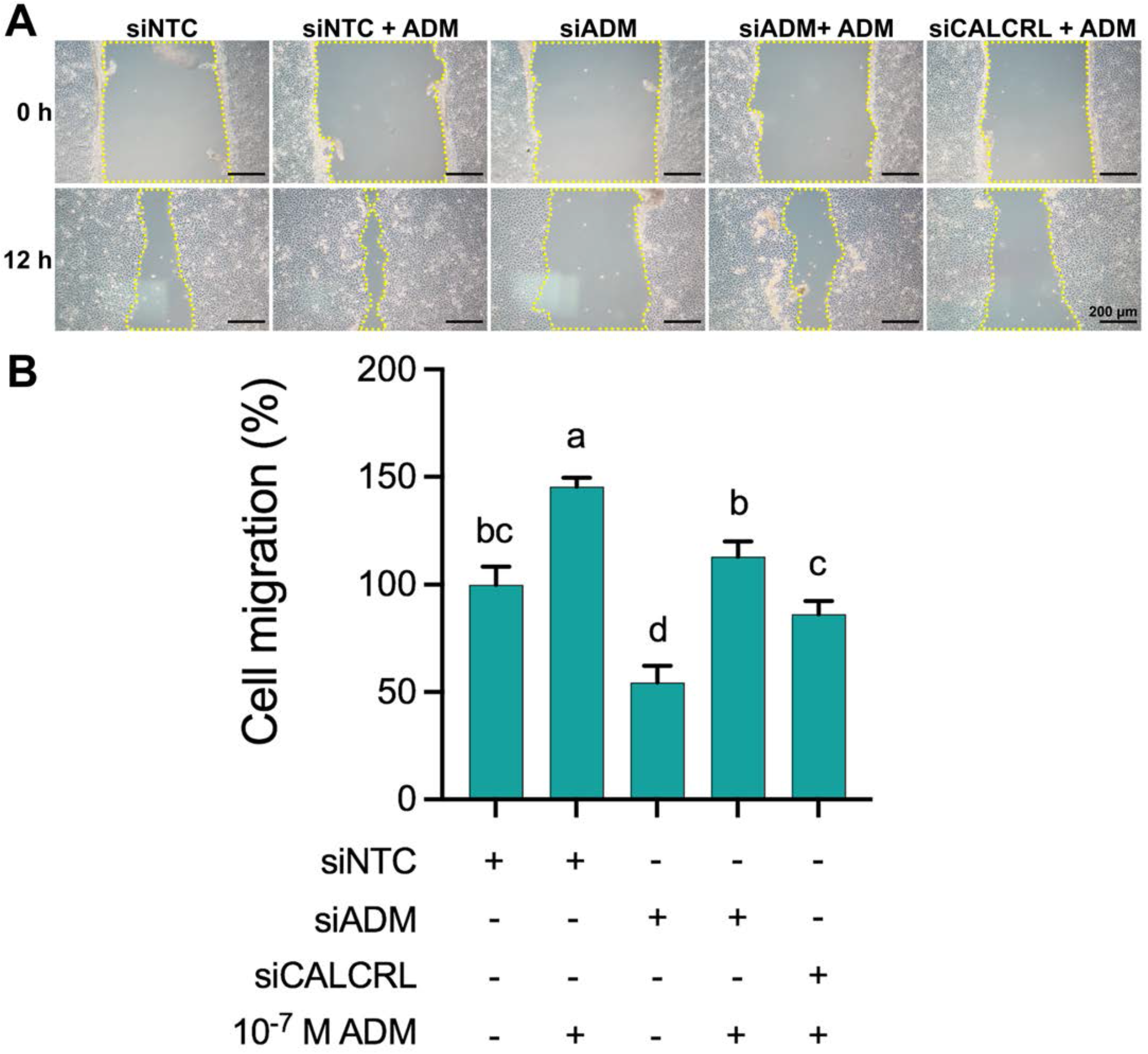
ADM stimulates migration of pTr1 cells via its receptor component CALCRL. The pTr1 cells (n=6 wells) were transfected with siNTC, siADM or siCALCRL for 48 h, and then subject to a 12-h wound-healing assay with or without 10^-7^ M ADM. Images (**A**) were taken immediately after scratching (0 h) and at 12 h post scratching to calculate area covered by cell migration (**B**). The dotted yellow lines indicate scratched borders of cell-free area. Bar=200 μm. Data are expressed as a percentage relative to siNTC treated cells. siNTC, nontargeting control siRNA; siADM, siRNA targeting porcine *ADM*; siCALCRL, siRNA targeting porcine *CALCRL*.Different superscript letters denote significant differences (*P*<0.05) among groups. Data are presented as means and SEM.

### ADM stimulates adhesion of pTr1 cells via its receptor component CALCRL

We then investigated the effects of ADM on attachment of pTr1 cells using siRNA-mediated knockdown (**Figure 3**). After 48 h transfection with siRNAs and 2 h attachment with or without exogenous ADM at 10^-7^ M, the number of adherent cells was increased (*P*<0.05) by 1.2-fold in both siNTC+ADM and siADM+ADM groups as compared to siNTC and siADM controls, respectively. Cell adhesion were not affected between siNTC and siADM groups regardless of the absence (siADM *versus* siNTC) and presence (siADM+ADM *versus* siNTC+ADM) of exogenous ADM. However, inhibition of CALCRL decreased (*P*<0.05) the ADM-derived attachment of pTr1 cells by 20% in siCALCRL+ADM as compared to siNTC+ADM.

**Figure. 3.**
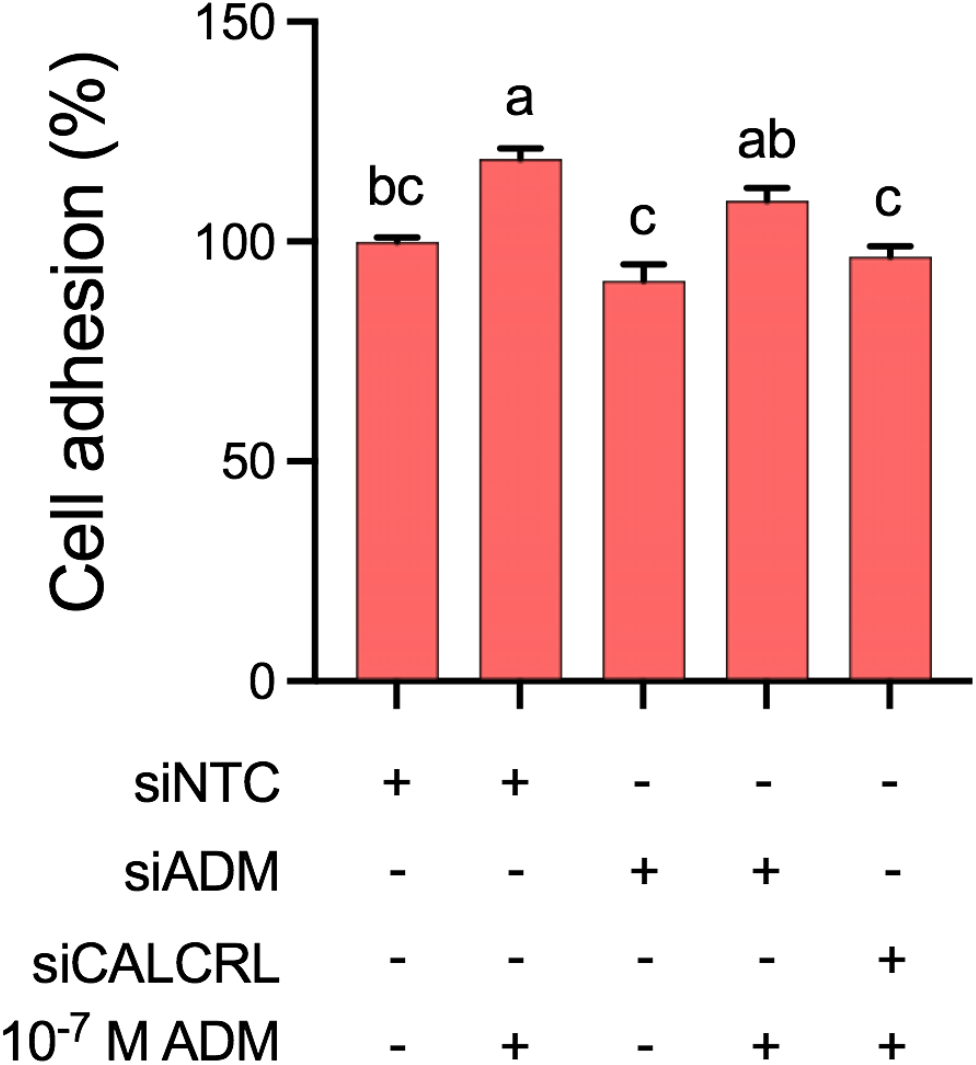
ADM stimulates adhesion of pTr1 cells via its receptor component CALCRL. The pTr1 cells (n=6 wells) were transfected with siNTC, siADM or siCALCRL for 48 h, and then re-seeded in 96-well plates with or without the presence of 10^-7^ M ADM. After 2 h for attachment, nonadherent cells were removed and adherent cells were determined. Data are expressed as a percentage relative to cells transfected with siNTC but no additional ADM treatment. siNTC, nontargeting control siRNA; siADM, siRNA targeting porcine *ADM*; siCALCRL, siRNA targeting porcine *CALCRL*. Different superscript letters denote significant differences (*P*<0.05) among groups. Data are presented as means and SEM.

### ADM stimulates migration and adhesion of pTr1 cells via activation of MTOR

Given that the MTOR inhibitor rapamycin (50 nM; (Wang et al. 2015a)) abrogated ADM-derived proliferative effects on pTr1 cells (Paudel et al. 2021), we then used rapamycin at the same dosage to evaluate whether the effects of ADM on migration and adhesion of pTr1 cells are also due to activation of MTOR (**Figures 4** and **5**). After 12 h incubation, exogenous ADM at 10^-7^ M increased (*P*<0.05) pTr1 cell migration by 1.3-fold in ADM group as compared to the nontreated control (**Figure 4**). This ADM-derived migration of pTr1 cells were abrogated (*P*<0.05) by 37% via inhibition of MTOR using rapamycin (rapamycin+ADM *versus* ADM). Interestingly, as compared to the nontreated control, rapamycin at 50 nM decreased (*P*<0.05) migration of pTr1 cells by 31% and 22% with or without exogenous ADM at 10^-7^ M, respectively. In addition, exogenous ADM at 10^-7^ M increased (*P*<0.05) the number of adherent pTr1 cells by 1.2-fold as compared to the nontreated control after 2 h incubation; whereas rapamycin reduced (*P*<0.05) such ADM-driven attachment of pTr1 cells by 25% (rapamycin+ADM *versus* ADM; **Figure 5**). Likewise, as compared to the nontreated control, rapamycin at 50 nM with or without exogenous ADM at 10^-7^ M decreased (*P*<0.05) the number of adherent pTr1 cells by 14%.

**Figure 4.**
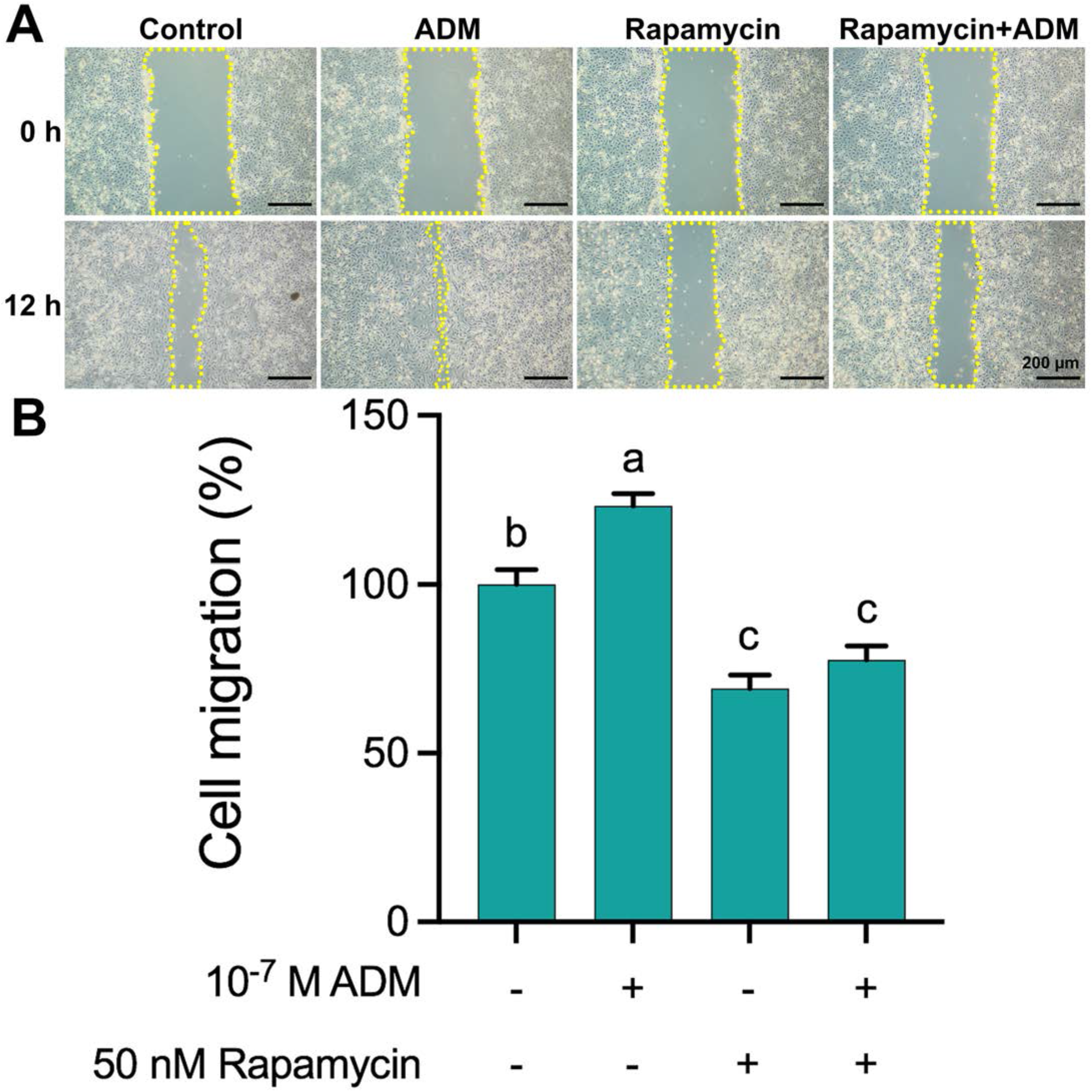
ADM stimulates migration of pTr1 cells via activation of MTOR. The pTr1 cells (n=6 wells) were pre-incubated with 50 nM rapamycin for 6 h, and then subject to a 12-h woundhealing assay with or without 10^-7^ M ADM. Images (**A**) were taken immediately after scratching (0 h) and at 12 h post scratching to calculate area covered by cell migration (**B**). The dotted yellow lines indicate scratched borders of cell-free area. Bar=200 μm. Data are expressed as a percentage relative to nontreated control cells. Different superscript letters denote significant differences (*P*<0.05) among groups. Data are presented as means and SEM.

**Figure 5.**
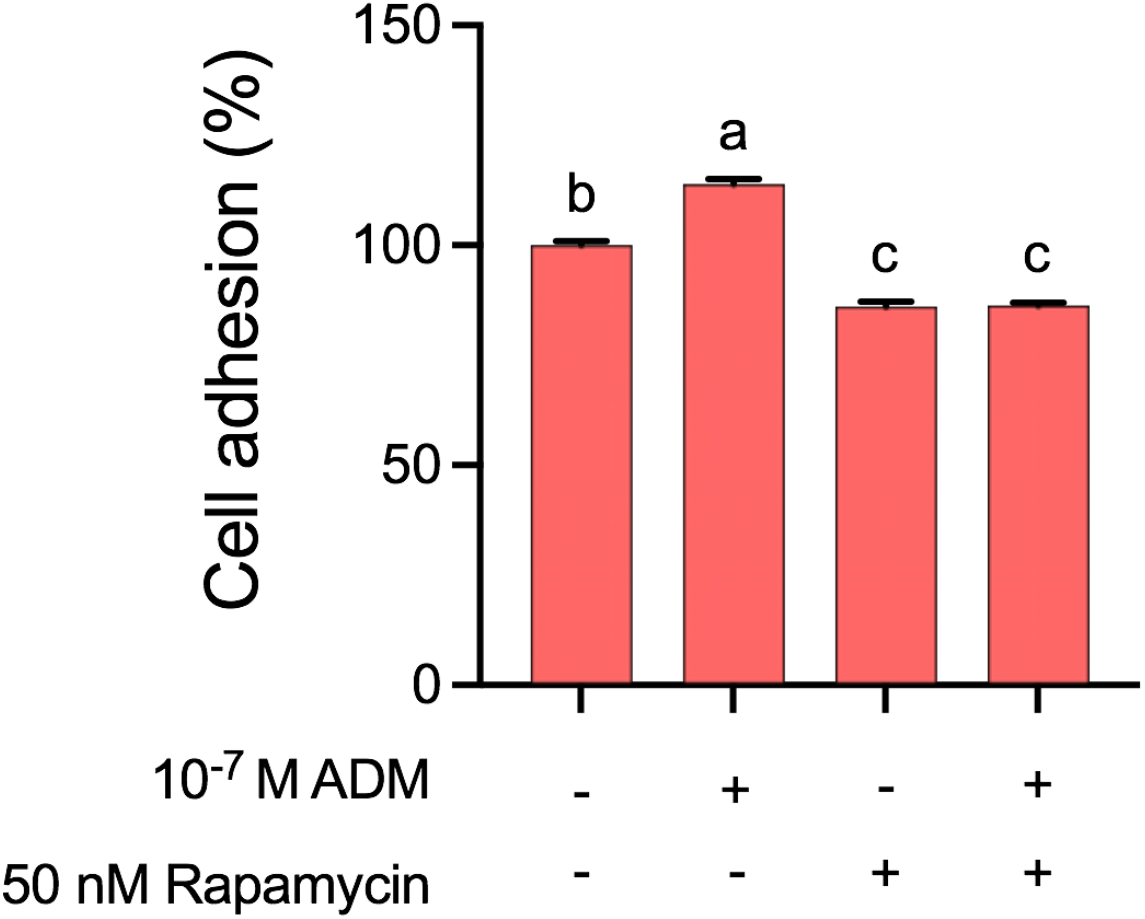
ADM stimulates adhesion of pTrl cells via activation of MTOR. The pTr1 cells (n=6 wells) were pre-incubated with 50 nM rapamycin for 6 h, and then re-seeded in 96-well plates with or without the presence of 10^-7^ M ADM. After 2 h for attachment, nonadherent cells were removed and adherent cells were determined. Data are expressed as a percentage relative to nontreated control cells. Different superscript letters denote significant differences (*P*<0.05) among groups. Data are presented as means and SEM.

### ADM activates AKT-TSC2-MTORC1 cell signaling pathway in pTr1 cells via its receptor component CALCRL

To further investigate whether and how ADM activates MTOR, particularly MTORC1 cell signaling pathway, we determined the protein expressions of total (t-) and phosphorylated (p-) MTOR, 4EBP1, P70S6K, S6, TSC2, and AKT in pTr1 cells at 30 and 60 min of ADM incubation using siRNA-mediated knockdown and Western blot analyses (**Figure 6**). After siRNA transfection and additional ADM treatment, the relative phosphorylation of MTOR (p-MTOR/t-MTOR) decreased (*P*<0.05) by 39% and 80% in siCALCRL-treated pTr1 cells as compared to siNTC at 30 and 60 min of ADM incubation, respectively (**Figure 6A**). As the direct downstream effectors of MTORC1, the relative phosphorylation of 4EBP1, P70S6K and S6 (target of P70S6K) were further determined (**Figures 6B-D**). The p-4EBP1/t-4EBP1 ratio decreased (*P*<0.05) by 27% and 56% in siCALCRL-treated pTr1 cells at 30 and 60 min of ADM incubation, respectively; but increased (*P*<0.05) by 1.5-fold in siNTC controls between 30 and 60 min of ADM incubation (**Figure 6B**). Likewise, the p-P70S6K/t-P70S6K ratio decreased (*P*<0.05) by 79% and 86% in siCALCRL-treated pTr1 cells at 30 and 60 min of ADM incubation, respectively; but increased (*P*<0.05) by 1.6-fold in siNTC controls between 30 and 60 min of incubation with ADM (**Figure 6C**). As the immediate target of P70S6K, the relative phosphorylation of S6 (p-S6/t-S6) decreased (*P*<0.05) by 39% and 43% in siCALCRL-treated pTr1 cells at 30 and 60 min of ADM incubation, respectively (**Figure 6D**). Moreover, we determined the relative phosphorylation of two upstream regulators of MTORC1, TSC2 and AKT (**Figure 6 E and F**). The p-TSC2/t-TSC2 ratio decreased (*P*<0.05) by 49% and 46% in siCALCRL-treated pTr1 cells at 30 and 60 min of ADM incubation, respectively (**Figure 6E**); whilst the p-AKT/t-AKT ratio decreased (*P*<0.05) by 79% and 89% (**Figure 6F**). In addition, ADM increased (*P*<0.05) both p-TSC2/t-TSC2 and p-AKT/t-AKT by 1.6- and 2.3-folds, respectively, in siNTC controls between 30 and 60 min of incubation (**Figure 6E** and **F**).

**Figure 6.**
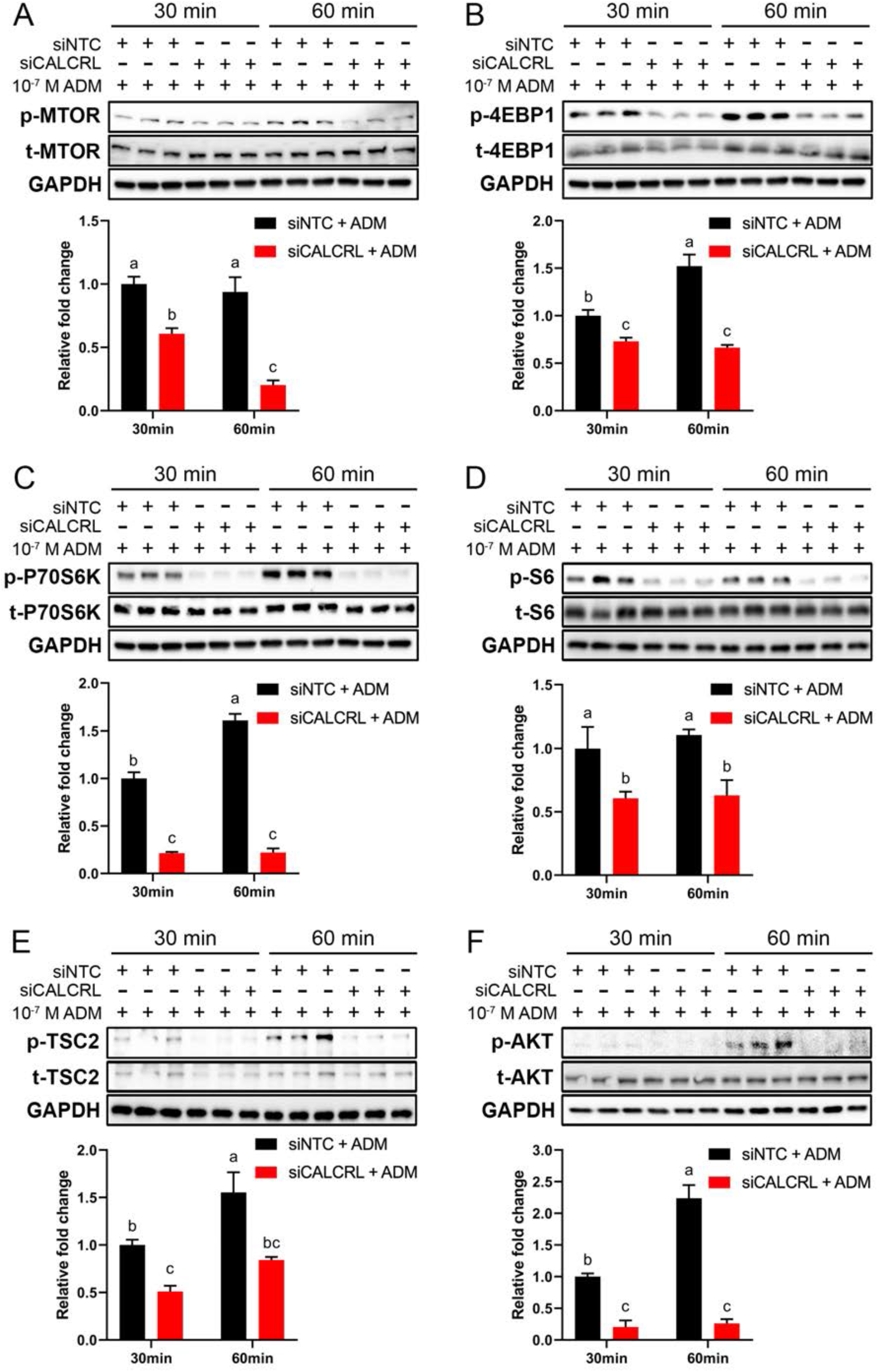
ADM activates AKT-TSC2-MTORC1 cell signaling pathway in pTr1 cells via its receptor component CALCRL. The pTr1 cells (n=3 wells) were seeded in 6-well plates transfected with siNTC or siCALCRL for 48 h before ADM treatment at 10^-^ M for additional 30- and 60-min. Images and quantification of Western blot analyses on total (t-) and phosphorylated (p-) MTOR (**A**), 4EBP1 (**B**), P70S6K (**C**), S6 (**D**), TSC2 (**E**), and AKT (**F**) in pTr1 cells. GAPDH was used as loading control antibody for each blot. siNTC, nontargeting control siRNA; siCALCRL, siRNA targeting porcine *CALCRL*. Data are expressed as relative fold change (p-to-t protein ratio). Different superscript letters denote significant differences (*P*<0.05) among groups.Data are presented as means and SEM.

## Discussion

In pigs, asynchrony between uterine signals and conceptus development limits surface area of placentation at the maternal-conceptus interface, leading to fetal crowding that eventually results in IUGR piglets at birth with poor survival in the neonatal period of life (Vallet et al. 2013; Bartol et al. 1993; Vallet et al. 2014; Wu et al. 2006). This represents a major constraint to improving reproductive performance in modern swine production enterprises as continuous selection for ovulation rate in sows and semen quality in boars have led to increased litter sizes from female with greater prolificacy (Johnson et al. 1999; Rothschild 1996; Tomes and Nielsen 1982). Therefore, it is imperative that functional roles of uterine secretions (i.e., histotroph) and underlying mechanisms regulating the growth and development of conceptuses in concert with intrauterine environment be established in pigs. Adrenomedullin (ADM) is a highly conserved peptide hormone required for intra-uterine spacing of blastocysts and angiogenesis during early pregnancy in rodents (Caron and Smithies 2001; Li et al. 2006; Matson et al. 2017; Lenhart and Caron 2012; Li et al. 2008). Our previous research demonstrated the temporal and spatial expression patterns of ADM and its receptors in the porcine uterus and peri-implantation conceptuses (Paudel et al. 2021). We also showed that ADM stimulates proliferation of pTr1 cells in a dose-dependent manner (10^-7^ M as the optimal dose tested) (Paudel et al. 2021). In this study, we further defined the functional impacts of ADM on proliferation, migration and adhesion of pTr1 cells, as well as the underlying mechanisms. Results of the present study significantly advanced understanding of ADM roles in support of the intricately orchestrated events required for elongation of the conceptus and establishment of a successful pregnancy in pigs.

Results of this study provide the first *in vitro* evidence that ADM acts on pTr1 cells to increase proliferation, migration and adhesion via activation of AKT-TSC2-MTORC1 cell signaling cascade. First, we confirmed that ADM increased pTr1 cell proliferation significantly (**Figure 1**), which is required for conceptus elongation in pigs and other ungulates (Kong et al. 2014; Wang et al. 2015a; Wang et al. 2015b; Bazer et al. 2015). These ADM-derived proliferative effects were abrogated by siRNA-mediated knockdown of either ADM or its shared receptor component CALCRL (**Figure 1**). Given that migration and attachment of conceptus Tr to LE are also prerequisite for implantation and establishment of a successful pregnancy (Bazer et al. 2015; Wang et al. 2016a; Erikson et al. 2009; Kim et al. 2010), we further determined the impacts of ADM on migration and adhesion of pTr1 cells. As a result, ADM significantly increased pTr1 cell migration (**Figure 2**) and adhesion (**Figure 3**), which were abrogated by siRNA-mediated knockdown of ADM or CALCRL. Interestingly, siRNA-mediated knockdown of ADM further decreased proliferative and migratory rates of pTr1 cells in siADM group as compared to siNTC control, suggesting the autocrine effects of endogenous ADM on pTr1 cell proliferation and migration (**Figures 1 and 2**). However, such reduction in the number of attached pTr1 cells was not observed in siADM group as compared to siNTC control, indicating that endogenous ADM *per se* may not be a stimulator for cell attachment. Instead, ADM may function like arginine (Arg), the functional amino acid that synergistically works with extracellular matrix proteins (e.g., SPP1) on pTr1 cells to facilitate the formation of focal adhesions, thereby increases in cell attachment (Wang et al. 2014b; Kim et al. 2010). Indeed, ADM signaling was among the top activated pathways triggered by Arg in both porcine and ovine Tr cells (**Paudel and Wang, unpublished data**). These results suggest that ADM acts on its receptor component CALCRL to stimulate proliferation, migration and adhesion of pTr1 cells.

MTOR is the master regulator of conceptus development in response to uterine histotroph during pregnancy in ungulates, including pigs, sheep and other ruminants (Wang et al. 2016c; Kong et al. 2014; Wang et al. 2015b, 2016a). As a serine/threonine kinase, MTOR is the catalytic subunit of two structurally distinct complexes: MTOR complex 1 (MTORC1) and MTOR complex 2 (MTORC2). MTORC1 includes MTOR, mammalian lethal with SEC13 protein 8 (MLST8) and regulatory-associated protein of MTOR (Raptor) and increases mRNA translation and cell proliferation (proliferative growth) (Wullschleger et al. 2006; Kong et al. 2014; Wang et al. 2015b). On the other hand, MTORC2 is composed of MTOR, MLST8 and rapamycin-insensitive companion of MTOR (Rictor) as the core components and is associated with cell migration, adhesion and cytoskeletal organization (spatial aspect of growth) (Wang et al. 2016a). Therefore, these effects of ADM on proliferation, migration and adhesion of pTr1 cells might be due to stimulation of MTOR, the shared component of MTORC1 and MTORC2. Our previous study revealed that ADM-induced proliferative effect on pTr1 cells was abrogated when MTOR activity was inhibited by rapamycin (Paudel et al. 2021). Here, we showed that rapamycin decreased both ADM-induced cell migration and attachment, demonstrating that ADM exerts both actions on pTr1 cells via activation of MTOR. In addition, cell migration and adhesion were further reduced in rapamycin-treated groups as compared to the nontreated control, suggesting that the stimulatory effect of endogenous ADM on migration and adhesion also be abrogated in pTr1 cells.

To understand how exogenous and endogenous ADM stimulates MTOR activity, we investigated effects of porcine ADM on activation of MTORC1 cell signaling cascade in pTr1 cells using siRNA-mediated knockdown of CALCRL. Once phosphorylated, MTOR becomes active member of MTORC1 that regulates cell proliferation in part by phosphorylating two best-characterized downstream effectors 4EBP1 and P70S6K, known regulators of protein synthesis (Hay and Sonenberg 2004). In the present study, ADM-induced phosphorylation of MTOR, 4EBP1, P70S6K and S6 (direct target of P70S6K) were decreased by siCALCRL, demonstrating that ADM-triggered effects on proliferation, migration and adhesion of pTr1 cells may be mediated by activation of MTOR activity, particularly MTORC1 signaling (**Figures 6A-D**). Meanwhile, TSC2 is a GTPase activating protein that stimulates GTPase activity of the small GTPase Rheb (Huang and Manning 2009; Inoki et al. 2003a; Huang and Manning 2008; Inoki et al. 2003b). Given that Rheb in its GTP bound form is an activator of MTORC1 (Inoki et al. 2003a), TSC2, in complex with TSC1, serve as an upstream regulator inhibitory to the MTORC1 signaling cascade, and phosphorylation of TSC2 removes that inhibition. Here, increases in p-TSC2/t-TSC2 ratio but abrogated by siCALCRL suggest that ADM bind to CALCRL to activate the MTORC1 signaling pathway by increasing phosphorylation of TSC2, thereby preserving Rheb in its GTP bound form in pTr1 cells (**Figure 6E**). Since inhibition of TSC2 activity usually results from AKT-mediated phosphorylation and membrane partitioning, we next investigated the abundances of total and phosphorylated AKT, a serine/threonine kinase that controls physiological processes, such as cell growth, survival and motility. In the present study, AKT phosphorylation was increased by ADM but decreased by siCALCRL, suggesting stimulatory effects of ADM on AKT-TSC2-MTOR, particularly MTORC1 signal transduction (**Figure 6F**). These findings help elucidate the mechanisms for functional roles of ADM in regulating embryonic survival, growth and development in pigs as well as other mammalian species.

In summary, this is the first report of functional roles of ADM in porcine conceptus trophectoderm cells *in vitro*. We have demonstrated that ADM binds to CALCRL to exert actions on proliferation, migration and adhesion of pTr1 cells. We also provided evidence that these stimulatory effects of ADM are regulated through activation of AKT-TSC2-MTOR, particularly MTORC1 signaling cascade. This work provides important insights into understanding the complexity of orchestrated events and underlying mechanisms responsible for conceptus-uterine signaling required for conceptus elongation, implantation and placentation. Moreover, it will guide the development of a novel strategy in using exogenous ADM or supplements to induce endogenous ADM in the uterine environment to prevent and treat IUGR in pigs and other mammals. Future studies will further dissect ADM actions on MTORC2 signaling transduction, cytoskeletal reorganization, as well as pre-implantation spacing of porcine conceptus Tr *in vitro* and *in vivo*. The regulatory mechanisms for ADM production in the uterine endometrium and conceptus Tr will also be investigated in pigs using organoid and explant culture.

## DECLARATIONS

### Conflict of interest

The authors declare that they have no conflict of interest.

### Ethical statement

An established cell line was used for this study. Thus, the present work did not require the approval for the use of animals by Institutional Animal Care and Use Committee of North Carolina State University.

### Informed consent

No informed consent is required for this study.

## ACKNOWLEDGEMENTS

Contributions of the graduate students and postdoctoral fellows from the Laboratory of Reproductive and Developmental Biology are gratefully acknowledged. This research was supported by Agriculture and Food Research Initiative competitive grant 2022-67015-36491 (to X.W.) from the U.S. Department of Agriculture National Institute of Food and Agriculture, and N.C. Agricultural Foundation grants 665143 and 665163 (to X.W.) from North Carolina State University.

## DATA AVAILABILITY

The data underlying this article will be shared on reasonable request to the corresponding author.

## AUTHOR CONTRIBUTION

X.W. conceived, designed the research and wrote the manuscript. B.L. performed the experiments and data curation. B.L. and S.P. contributed to experimental design and data analysis. W.F. and J.P. contributed to manuscript editing. All authors reviewed the manuscript.

## Notes

**Grant Support:** This research was supported by Agriculture and Food Research Initiative competitive grant 2022-67015-36491 (to X.W.) from the U.S. Department of Agriculture National Institute of Food and Agriculture, and N.C. Agricultural Foundation grants 665143 and 665163 (to X.W.) from North Carolina State University

### Competing Interest Statement

The authors have declared no competing interest.

